# Prediction of neural activity in connectome-constrained recurrent networks

**DOI:** 10.1101/2024.02.22.581667

**Authors:** Manuel Beiran, Ashok Litwin-Kumar

## Abstract

We develop a theory of connectome-constrained neural networks in which a “student” network is trained to reproduce the activity of a ground-truth “teacher,” representing a neural system for which a connectome is available. Unlike standard paradigms with unconstrained connectivity, here the two networks have the same connectivity but different biophysical parameters, reflecting uncertainty in neuronal and synaptic properties. We find that a connectome is often insufficient to constrain the dynamics of networks that perform a specific task, illustrating the difficulty of inferring function from connectivity alone. However, recordings from a small subset of neurons can remove this degeneracy, producing dynamics in the student that agree with the teacher. Our theory can also prioritize which neurons to record from to most efficiently predict unmeasured network activity. Our analysis shows that the solution spaces of connectome-constrained and unconstrained models are qualitatively different and provides a framework to determine when such models yield consistent dynamics.

## INTRODUCTION

Establishing links between the connectivity of large neural networks and their emergent dynamics is a major goal of theoretical neuroscience. Many studies have attempted to develop methods to infer synaptic connectivity from functional correlations derived from recorded neural activity. However, this “inverse problem” has proven to be challenging and often ill-posed ^1–5^, due to the degeneracy of the space of network connectivities that produce similar dynamics. Such inference is particularly difficult when neural dynamics are low-dimensional or otherwise structured ^1^.

The recent availability of comprehensive synaptic connectome datasets has led to approaches that focus instead on the “forward problem” of predicting neural dynamics and function from synaptic connectivity. The scale of such datasets has increased rapidly, from the 302 neurons of the nematode *C. elegans* identified decades ago^6^ to recently acquired volumes containing entire nervous systems of *Drosophila* larvae^7^ and adults^8–10^, larval zebrafish^11^, and other model systems. Several studies have compared connectomes with functional connectivity based on activity correlations between neurons in resting state or in response to optogenetic perturbations^12^. This has highlighted striking differences for certain systems^13^. A complementary line of research has made use of connectome information to initialize or build explicit priors on the distribution of the parameters of neural network models ^14;15^. In some cases, these models are then trained with machine learning algorithms to perform computations, and it has been found empirically that such biological constraints sometimes yield models with better generalization properties or ability to predict neural data^16–19^. However, the ill-posedness of the inverse problem and lack of one-to-one correspondence between structure and function call into question whether such approaches can yield consistent predictions.

A major challenge for connectome-constrained models is uncertainty in synaptic or neuronal biophysical parameters that affect neural dynamics. Connectomes generated from electron-microscopy imaging provide information on structural connections, neurotransmitter identities of chemical synapses^10;20^, and connection strengths estimated by synapse count^21^ or volume ^22^. However, other biological processes are undetermined, such as the neuromodulatory environment, existence of electrical synapses, and functional properties of individual neurons^23^. Changes in such parameters have previously been shown to produce dramatic alterations in network activity ^24–27^.

In this study, we develop a theory of the solution spaces of networks with specified connectivity but unknown single-neuron biophysical parameters. These parameters account for unmeasured factors that drive neural activity and the heterogeneity of functional properties across neurons^28;29^. We use a “teacher-student” paradigm in which the activity of a “student” network is trained to reproduce the activity of a “teacher” network. The teacher is a synthetic model that represents ground truth, analagous to a neural system for which a connectome is available. Unlike previous theories in which the student and teacher neurons have the same input-output function and the connectivity is trained^1;30^, here we assume that the teacher and student connectivities are the same but the biophysical properties of the student neurons are a priori different from those of the teacher. These undetermined parameters represent uncertainty and model mismatch between student and teacher.

We find that training a connectome-constrained student neural network to generate the task-related readout of the teacher is not enough to produce consistent dynamics in the teacher and student. Multiple combinations of single neuron parameters, each producing different activity patterns, can equivalently solve the same task. However, when connectivity constraints are combined with recordings of the activity of a subset of neurons, it is possible to predict the activity of neurons that have not been recorded. The minimum number of recordings depends on the dimensionality of the network dynamics, rather than the total number of neurons. This contrasts with student networks whose connectivity is unconstrained, which always display degenerate solutions. Interestingly, even when neural activity is well-reconstructed, specific values of unknown single-neuron parameters are often not recovered accurately, suggesting that some combinations of parameters are “stiff,” with strong effects on neural dynamics, while other ones are “sloppy,” not having a strong effect. Our theory can also rank neurons that should be recorded with higher priority to maximally reduce uncertainty in network activity, suggesting approaches that iteratively refine network models using neural recordings.

## RESULTS

### Teacher-student recurrent networks

To explore how a connectome constrains the solutions of neural network models, we studied a teacher-student paradigm^31;32^. A recurrent neural network (RNN) that we call the teacher is constructed, and the parameters of a student RNN are adjusted to mimic the teacher. In this work, the teacher is used as a proxy for a neural system whose connectome has been mapped. To describe both teacher and student networks, we use RNNs composed of *N* firing rate neurons, in which the activity of neuron *i* is described by a continuous variable *r*_*i*_ (*t*) (see Methods for details). The activity is a nonlinear function of the input current that the neuron receives, *x*_*i*_ (*t*), and depends on the single neuron parameters. For instance, if we describe the input-output function using parameters for each neuron’s gain and bias, the activity of neuron *i* is

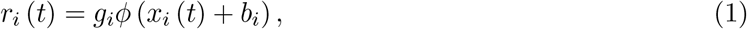

where *g*_*i*_ and *b*_*i*_ are the gain and bias parameters of the *i*th neuron, and *ϕ* is a nonlinear function. The dynamics are then given by:

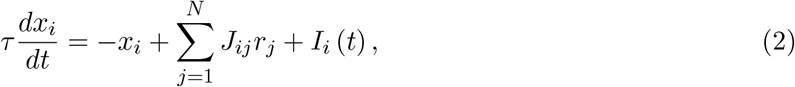

where *J*_*ij*_ corresponds to the synaptic weight from neuron *j* to neuron *i*, and *I*_*i*_ (*t*) is the time-varying external input received by neuron *i*. Note that the number of unknown parameters in the student network scales differently depending on whether single-neuron parameters or connectivity parameters are free. There are *N* ^2^ free synaptic strength parameters if the connectivity is unspecified, as in conventional machine learning models and prior studies of teacher-student paradigms^31;32^. On the other hand, for connectome-constrained networks, the number of unknown parameters is proportional to *N*. For example, when we parameterize the input-output function of neurons with gains and biases, as in Equation (1), there are 2*N* unknowns.

### Student network constrained by task output

We first asked whether teacher and student networks that share the same connectivity matrix *J* exhibit consistent solutions when the student is trained to reproduce a task performed by the teacher (Fig. 1). Because we are interested in whether connectivity constraints yield mechanistic models of the teacher’s behavior, we measure the consistency of solutions using the similarity of the activity of neurons in the teacher and those same neurons in the student. Such a direct comparison is possible because the connectome uniquely identifies each individual neuron. We also measure the similarity of teacher and student single neuron parameters. We refer to the dissimilarity between teacher and student activities or parameters as the “error” associated with each respective quantity. We note that our notion of similarity between teacher and student is more precise than requiring similarity of collective dynamics as measured through dimensionality reduction methods, such as principal components analysis. Indeed, matching such dynamics can be accomplished by recording a small number of neurons without access to a connectome^33;34^.

**Figure 1:**
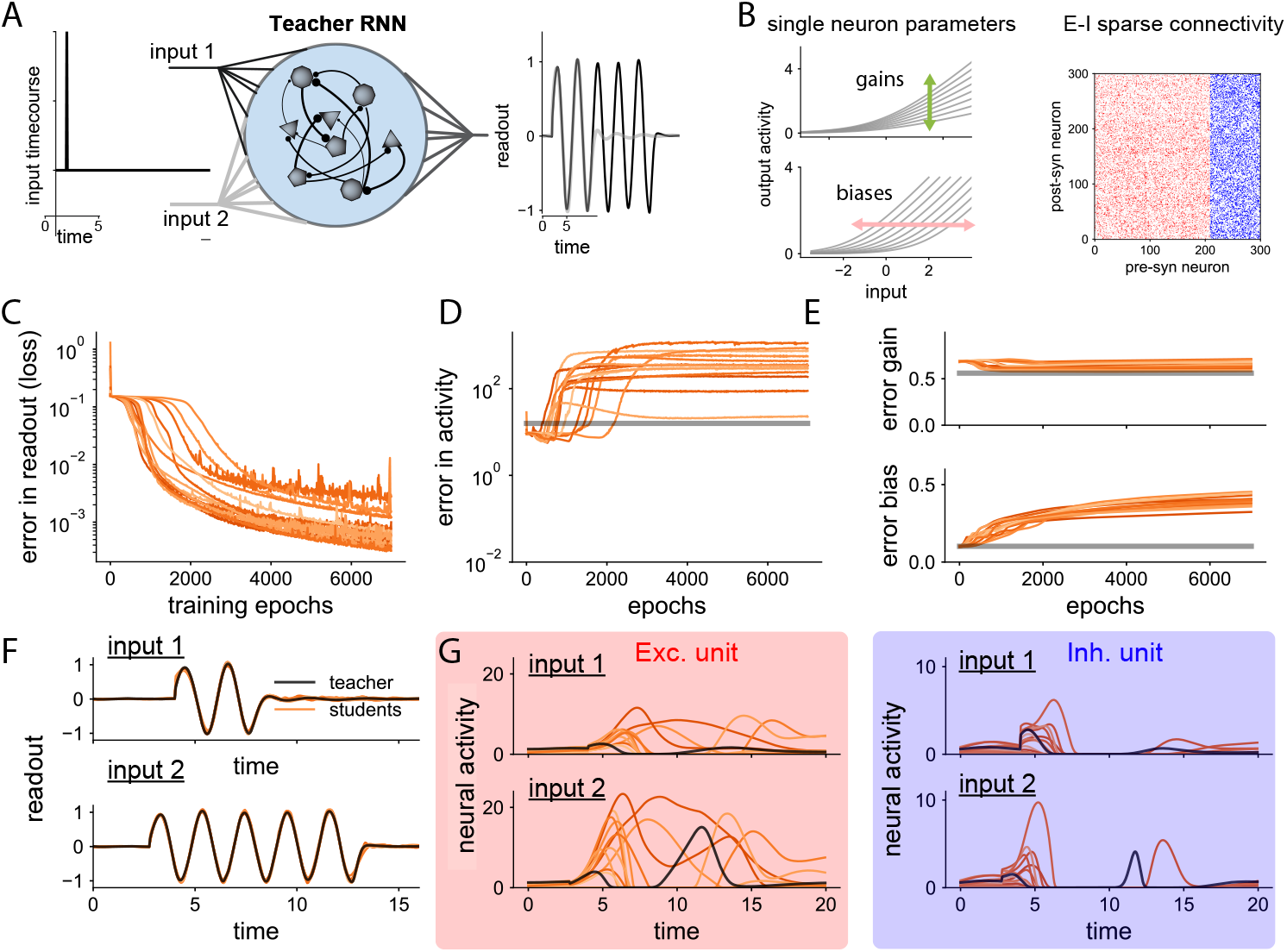
Task-constrained networks with the same connectivity. **A** A teacher RNN is trained to generate two different readout responses based on two different output pulses. **B** Properties of the teacher RNN. The teacher RNN has heterogeneous single neuron parameters (gains and biases of single neuron activation functions, left), sparse connectivity with connection probability *p* = 0.5 (right), and neurons connect through either excitatory (red) or inhibitory (blue) synapses. **C** Student networks with the same connectivity as the teacher are trained to produce the teacher’s output. Error in the readout (training loss, MSE) as a function of training epochs. Each colored line corresponds to a different student network. **D** Error (mismatch in neural activity) between teacher and student RNNs. Gray line, for reference, corresponds to the average error in activity when the student reproduces the teacher’s activity but with shuffled neuron identities. **E** Error in gains and biases vs. training epochs. **F** Readout of teacher and student networks after training, for the two trial types (top and bottom). Teacher and student networks both solve the task. **G** Neural activity of an example excitatory (left) and inhibitory (right) neuron. Teacher and student neurons exhibit different single-neuron dynamics.

We built a teacher network that is able to perform a flexible sensorimotor task. Specifically, the network implements a variant of the cycling task^35^, which requires oscillatory responses of different durations to be produced, based on a transient sensory cue (Fig. 1 A; see Methods). In the model, firing rates are a non-negative smooth function of the input currents, and the unknown single neuron parameters are the gains and biases, representing heterogeneity in individual neurons’ excitabilities and thresholds (Fig. 1 B left). The connectivity matrix is sparse and obeys Dale’s law, so that neurons are classified as excitatory or inhibitory (Fig. 1 B right).

We trained multiple students to generate the same readout signals as the teacher. Each student is initialized with different gains and biases before being trained via gradient descent. Trained networks successfully reproduce the teacher’s readout (Fig. 1 C, F). However, the error in the neural activity of the student, compared to the teacher, increases over training epochs (Fig. 1 D). As a baseline, we computed the error of a student whose neurons match the activities of all neurons in the teacher, but with shuffled identities (gray line, Fig. 1 D). In this baseline, the manifold of neural activity is the same in teacher and student, but not the activity at the level of single neurons. In all networks, the error in activity after training remains above this baseline, indicating that training does not produce a correspondence between the function of individual teacher and student neurons. Fig. 1 G shows the activity of one example excitatory and one inhibitory neuron, across different students and the teacher. The neuronal dynamics across different student networks is highly variable, and all students differ from the teacher.

Finally, we examined the error in single neuron parameters, i.e. the gains and biases, between teacher and student (Fig. 1 E). The error in gains does not vary much over training and is comparable to a randomly shuffled baseline. The error in biases grows slightly, although it remains within the same order of magnitude as the shuffled baseline.

We conclude that, in general, knowledge of synaptic connectivity and task output in recurrent networks is not enough to predict the activity of single neurons. For the task we considered, there is a degenerate space of network solutions, given by different combinations of single neuron gains and biases, that generate different patterns of recurrent neural activity whose readout solves the same task. We note that there may be scenarios for which this degeneracy is reduced (see Supp. Figs. 8, 9 and Discussion), such as small networks optimized for highly specific functions. In these cases, random choices of single neuron parameters may lead to above-chance prediction of neural activity (Supp. Fig. 9). Nonetheless, our results show that, even with *N* ^2^ connectivity constraints, task-optimized neural dynamics are, in general, highly heterogeneous.

### Student network constrained by activity recordings

We next asked whether these conclusions change if, instead of recording only the task-related readout activity, we record the activity of a subset of neurons in the teacher network. We use *M* ≤ *N* to denote the number of recorded neurons. The student is trained to reproduce this recorded activity, which provides additional constraints on the set of possible students (Fig. 2 A,B). The recording of subsampled activity in the teacher is analogous to neural recordings in imaging or electrophysiology studies, where only a subset of neurons are registered. We trained two types of student networks: students that have access to the teacher connectome, and students that are not constrained in connectivity. For connectome-constrained students (Fig. 2 C,E), the neuronal parameters of both recorded and unrecorded neurons are unknown and therefore trained. For students with unconstrained connectivity, the connectivity parameters are trained instead. To provide a fair comparison, in those cases we set the neuronal parameters of the student equal to those of the teacher (Fig. 2 D,F). Additionally, given that there is not a direct map between unrecorded neurons in the teacher and the student when the connectome is not known, after training the connectivity, we search for the mapping between student and teacher neurons that minimizes the mismatch in unrecorded activity using a greedy algorithm (see Methods).

**Figure 2:**
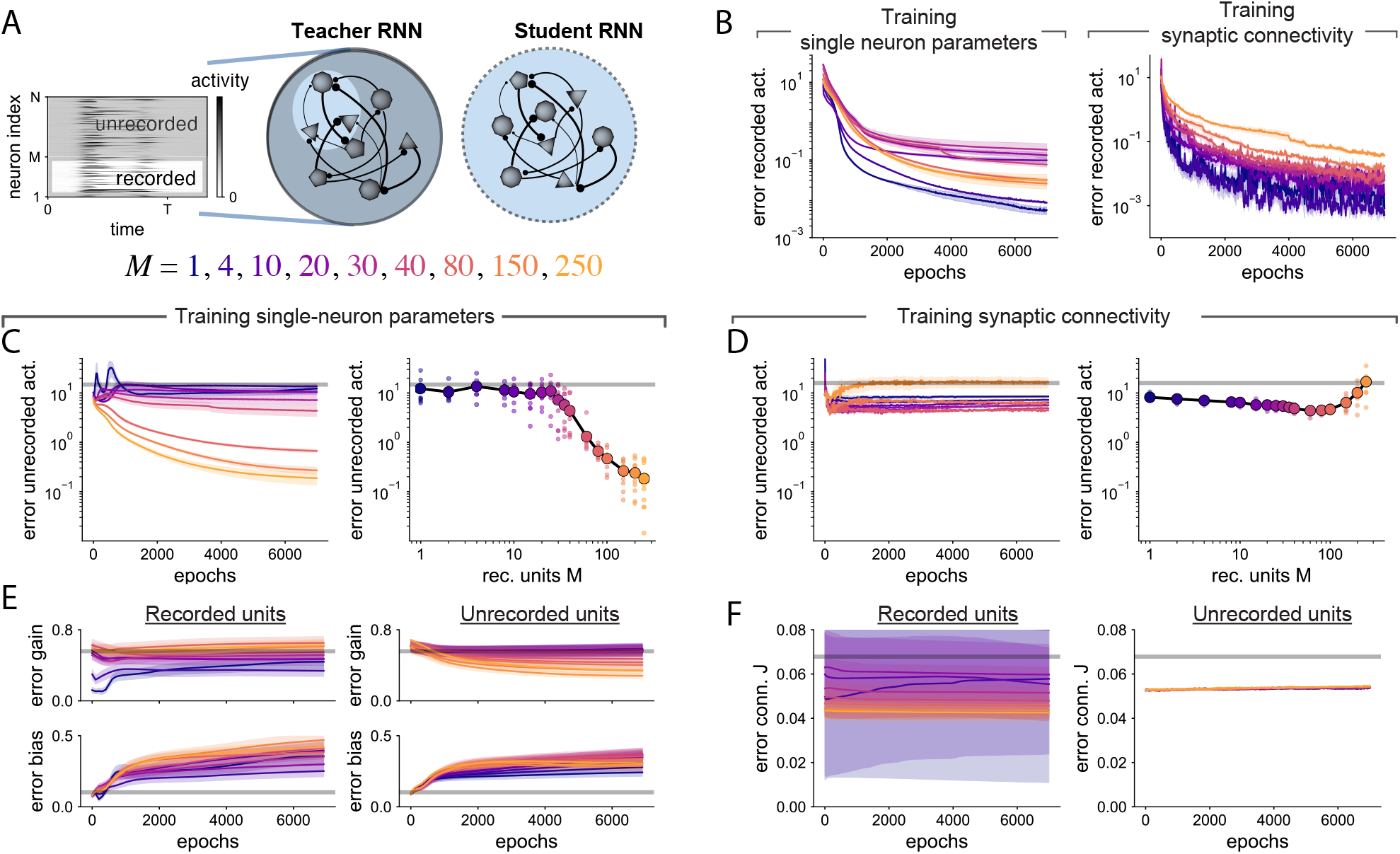
Transition in the prediction of the activity of unrecorded neurons when training singleneuron parameters. **A** The student RNN is trained to mimic the activity of *M* recorded neurons in a teacher RNN. **B** Error in recorded activity (loss) vs. training epochs for students with trained single neuron parameters (left) and students with trained connectivity (right). Lines correspond to different numbers of recorded neurons *M* and show mean and SEM over 10 random seeds. All students successfully reproduce the recorded activity of the teacher after learning. **C** Left: Error between teacher and students in the activity of the *N* − *M* unrecorded neurons vs. training epochs. Right: Error in unrecorded neuronal activity after training, as a function of number of recorded neurons *M*. Error is substantially reduced when recording from *M >* 30 neurons. **D** Analogous to Panel D, but training synaptic weights instead. The error in the activity of unrecorded neurons remains high across values of *M*. **E** Error in gains and biases vs. training epochs. Left: Parameters of recorded neurons. Right: Parameters of unrecorded neurons. **F** Analogous to panel E for connectivity weights between recorded neurons (left) and between unrecorded neurons (right).

We found that both connectome-constrained and unconstrained students are able to mimic the activity of the *M* recorded neurons in the teacher (Fig. 2 B, the teacher has *N* = 300 neurons). Although there are small differences for different values of *M*, all students achieve small errors. We then asked whether the activities of unrecorded neurons are consistent between teacher and student. When the connectivity is provided (Fig. 2 C), the error for unrecorded neurons is dramatically reduced when more than *M*^∗^ = 30 neurons in the teacher are recorded (example in Supplementary Fig. 11). Thus, when recording from enough neurons, the connectome provides sufficient information to also recover the activity of the remaining unrecorded neurons. In comparison, when training the synaptic weights (Fig. 2 D), unrecorded neurons’ activities are not recovered substantially better than baseline (see Methods for pairing of unrecorded neurons between teacher and student). These networks therefore exhibit degenerate solution spaces even when a majority of neurons in the network are recorded.

We then assessed whether the neuronal parameters of student networks converge to those of the teacher after training. For students whose connectivity is trained, the error in synaptic weights remains high, both for connections between recorded and unrecorded neurons (Fig. 2 F). We may expect this to occur given that the activity of unknown neurons in these networks is not well-predicted (Fig. 2 D). More surprisingly, errors in single neuron parameters in connectome-constrained networks also remain high, even when the activity of unknown neurons is well-predicted (Fig. 2 E). In particular, errors in biases increase over training, while errors in gains are only slightly reduced. We did not find qualitative differences in the behavior of single neuron parameters for recorded and unrecorded neurons.

So far, we focused on neural activity that is generated by recurrent interactions, triggered by brief external pulses. We further explored whether similar results hold when the teacher and student are driven by a time-varying external input (Fig. 8). Additionally, we systematically varied the distributions of gains distributions and connectivity sparsity in input-driven networks (Supp. Fig. 9). For these networks, the prediction of unrecorded neural activity prior to training, as compared to a randomly shuffled baseline, exhibited variability, with certain networks exhibiting above-chance prediction accuracy even with random parameters. Nonetheless, the qualitative dependence of prediction accuracy on number of recorded neurons was unchanged. These results suggest that, while certain features of neural activity may be predictable even with random parameters, improving upon this baseline through training requires sufficiently many recorded neurons.

In summary, connectome-constrained networks are able to predict the activity of unrecorded neurons when constrained by the activity of enough recorded neurons. In contrast, networks with trained connectivity cannot predict unrecorded activity. Nevertheless, in all cases, the unknown parameters are never precisely recovered, suggesting that multiple sets of biophysical parameters can lead to the same neural activity.

### Required number of recorded neurons is independent of network size

We have shown that recording from a subset of neurons in an RNN performing a task can yield consistent predictions about unrecorded neural activity, when a connectome is available. What features of the RNN determine the required number of recorded neurons? We considered two alternatives: the required number is a fixed fraction of the total number of neurons in the network, or the number is determined by properties of the network dynamics rather than network size. The former alternative would pose a challenge for large connectome datasets.

To disambiguate these two possibilities, we examined a class of teacher networks whose population dynamics are largely independent of their size. We generated networks with specific rank-two connectivity that autonomously generate a stable limit cycle ^36^ (Fig. 3 A and Methods). In these networks, the input currents received by each neuron oscillate within a two-dimensional linear subspace, independent of network size (Fig. 3 B).

**Figure 3:**
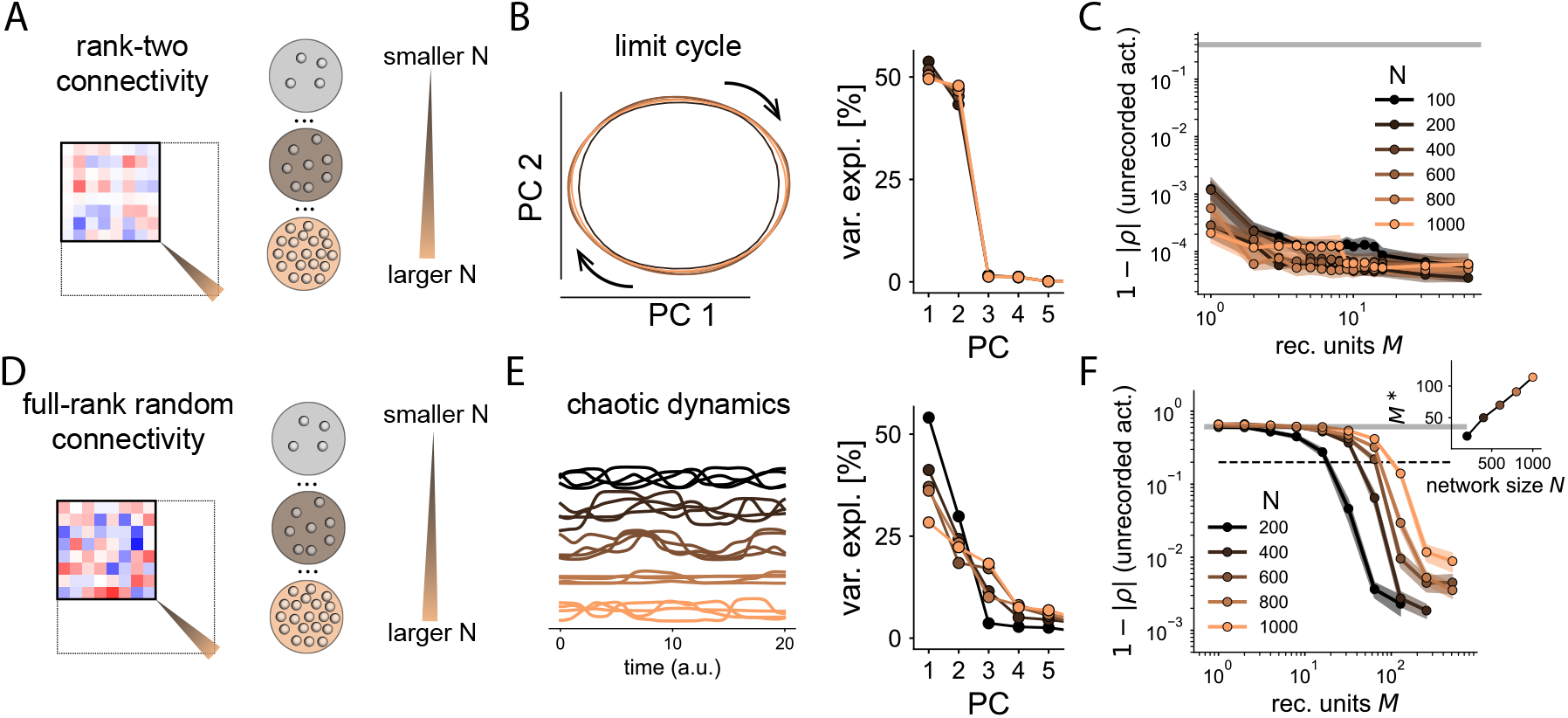
Prediction of unrecorded neurons’ activity depends on the geometry of dynamics, not network size. **A** Set of teacher RNNs with variable network size *N* but fixed rank of the connectivity matrix (see Methods). **B** Teachers of different size produce the same low-dimensional dynamics. Left: Dynamics projected on the top two principal components (PCs). All RNNs generate a limit cycle largely constrained to a 2D linear subspace. Right: Variance screeplot. **C** Error in the activity of unrecorded neurons, after training. We measured the correlation distance between activity in the teacher and student. Empirical average and SEM for each network size (10 networks per condition). Gray line sets represents the baseline error corresponding to shuffled neuronal identities. **D** Set of teacher RNNs with variable network size *N* with random (full-rank) connectivity. **E** Left: Networks generate high-dimensional chaotic dynamics. Sample activity of four units for networks of different sizes. Right: Variance scree plot. Larger networks generate higher dimensional dynamics. **F** Error in the activity of unrecorded neurons after training. Larger networks require recording from a larger number of neurons *M* to predict unrecorded activity. Inset: Number of neurons *M*^∗^ needed to predict unrecorded activity above a certain threshold (set to 0.2; dotted line), as a function of network size.

We found no difference in a plot of error in unrecorded activity against number of recorded neurons *M*, for networks of different sizes (Fig. 3 C), suggesting that accurate predictions can be made when recording from few neurons, even in large networks. Examining more closely the dependence of the error on the number of recorded neurons *M* for these networks, we observed that when *M* = 1, the error is substantially higher than when *M* = 7. In the former case, the student network produces oscillatory activity with the same frequency as the teacher, but the activity of unrecorded neurons exhibits consistent errors at particular phases of the oscillation (example in Supplementary Fig. 10). In contrast, when *M* = 7, errors in recorded and unrecorded neurons are comparable.

These results led us to hypothesize that the number of recorded neurons required to accurately predict neural activity scales with the dimensionality of the neural dynamics, not the network size. This would explain why networks with widely varying sizes but similar two-dimensional dynamics exhibit comparable performance (Fig. 3 C).

To further test this hypothesis, we studied a different setting in which we trained students to mimic another class of teacher networks, strongly-coupled random networks ^37^ (Fig. 3 D). In such networks, the neural activity is chaotic and, in contrast to the low-rank oscillating networks (Fig. 3 B), the linear dimensionality of the dynamics grows in proportion to the number of neurons^38^ (Fig. 3 E). In this case, we found that the number of recorded neurons *M*^∗^ required to predicted the activity of unrecorded neurons grows proportionally with the network size (Fig. 3 F). In total, these results suggest that recording from a subset of neurons, on the order of the dimensionality of network activity, is sufficient to improve prediction of unrecorded neural activity. Later, we will show that this numerical result is consistent with the predictions of an analytical theory.

### Robustness to model mismatch

So far, we have considered teacher and student networks that belong to the same model class of firing-rate networks with parameterized input-output functions and connectivity. However, when building models based on experimental data, there will always be some degree of “model mismatch” due to unaccounted or incorrectly parameterized biophysical processes. Moreover, errors in synaptic reconstruction and inter-individual variability in connectomes imply that connectivity estimates will also be associated with imprecision^39^. In this section, we examine whether our qualitative results hold when the student is described with a different model class than the teacher.

We used the same teacher network as in Figs. 1 And 2. We focused on two types of differences in the structure of student network (Fig. 4 A-B). In one case, we used a different activation function in the student network and the teacher (Fig. 4 B). Up to considerable amounts of change in the activation functions, there is a steep decrease in the error of unrecorded neurons as the number of recorded neurons is increased (Fig. 4 C right), similar to the case without model mismatch. However, there are small differences in the error of the recorded activity depending on the mismatch. In particular, the error in recorded activity is higher when the activation function is flatter than that of the teacher (Fig. 4 C left), although in all cases the error in recorded neurons after training is reduced by at least an order of magnitude. As the mismatch in activation functions increases, it becomes harder to exactly reproduce the recorded activity.

**Figure 4:**
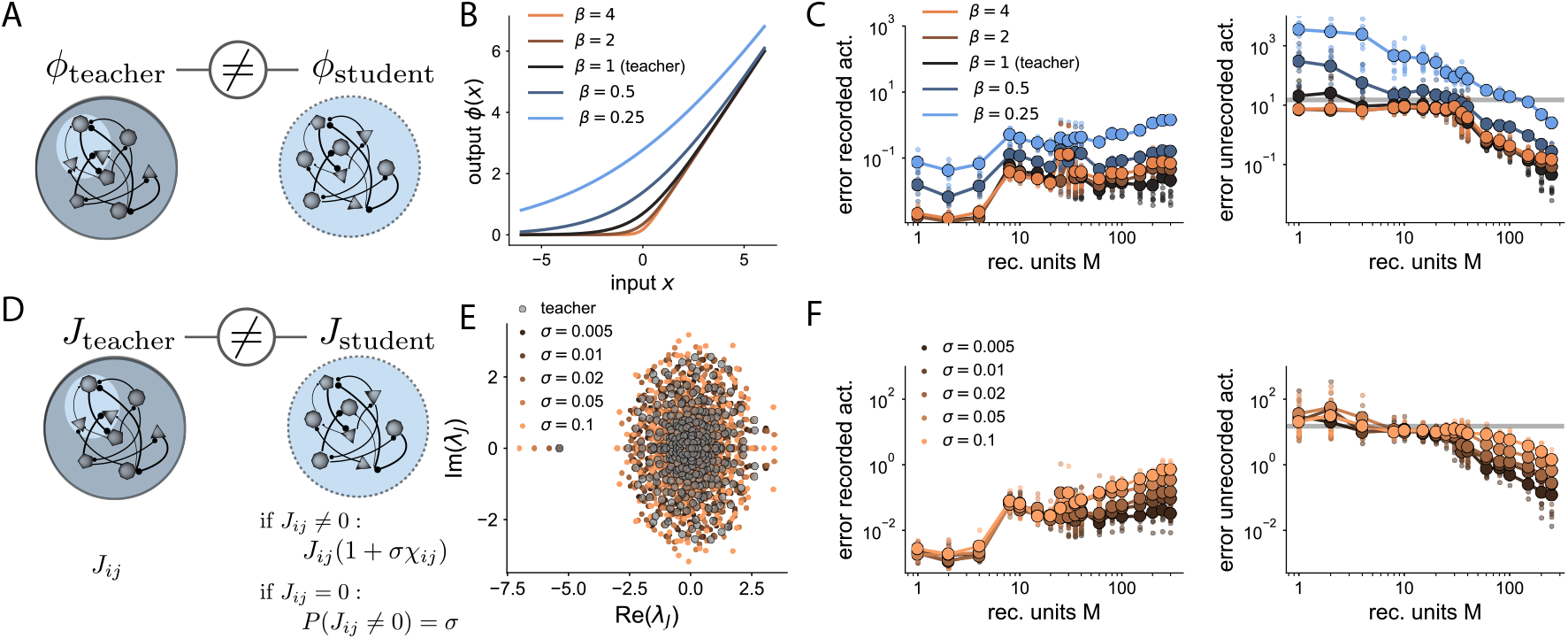
Model mismatch between teacher and student. **A** Mismatch in activation functions of teacher and student neurons. **B** The activation function is a smooth rectification but with different degrees of smoothness, parameterized by a parameter *β*. Teacher RNN from Fig. 2. **C** Errors in the activity of recorded (left) and unrecorded (right) neurons for different values of model mismatch between teacher and student. Within a large range of mismatch we observe a decrease in the error in unrecorded neurons when *M >* 30. **D** Mismatch in the connectivity between teacher and student, mimicking errors in connectome reconstruction. **E** Eigenvalues of the teacher and student connectivity matrices, for different levels of connectivity mismatch. **F** Errors in the activity of recorded (left) and unrecorded (right) neurons for different levels of mismatch in the connectivity.

We then considered errors in the measured connectivity of the student network. We simulated such errors using two different random processes (Fig. 4 D): for existing connections, we added Gaussian noise to the connection strength, while for non-existent connections, we added spurious connections with some probability *σ* (see Methods). We then used this corrupted version of the connectivity in the student network. Adding noise to the connectivity shifts all eigenvalues of the connectivity spectrum (Fig. 4 E) and modifies their corresponding eigenvectors. When the single-neuron parameters of the student are trained to match the recorded activity of the teacher, both the error in recorded and unrecorded activity increase smoothly as connectivity noise is increased (Fig. 4 F). We also again found a steep decrease in the prediction of unrecorded neurons as a function of *M*, suggesting that this qualitative behavior is not overly sensitive to deviations from the true connectivity.

Can model mismatch due to single-neuron parameters be compensated by allowing allowing the connectivity to be trained? To examine this, we trained the connectivity of the student instead of its gains and biases in a scenario with model mismatch in the activation functions (Supp. Fig. 11). The student network is initialized with the same connectivity as the teacher, before the synaptic weights are trained. We found that the activity of unrecorded neurons is less well-predicted when the connectivity, rather than single neuron parameters, is trained. We conclude that allowing for changes in single neuron parameters instead of connectivity proves to be more effective to account for the mismatch in the activation functions, illustrating the importance of parameterizing this particular form of uncertainty.

The numerical experiments in this section show that predictions about unrecorded neural activity can be made in connectome-constrained networks even when there is a model mismatch between student and teacher. These results do not imply that simplified models such as those we have studied here will be sufficient to make accurate predictions in all settings. However, they suggest that some degree of mismatch is tolerable (see Discussion).

### Linear network model

To analytically describe the properties of connectome-constrained networks, we developed a theory of our teacher-student paradigm. The theory aims to explain, first, how the teacher and student produce the same activity despite different single neuron parameters, and second, the conditions under which the student’s activity converges to that of the teacher.

We begin with a simplified linear model and, in subsequent sections, relax assumptions and show that the main insights remain valid. In the simplified model, the teacher and student RNNs have linear single-neuron activation functions, the only unknown single neuron parameters are the biases *b*_*i*_, and the connectivity *J* is a matrix with rank *D* (Fig. 5 A). This rank constraint implies that recurrent neural activity is confined to a *D*-dimensional subspace of the *N* -dimensional neural activity space. We focus on the network’s steady-state activity at equilibrium, which depends linearly on the biases:

**Figure 5:**
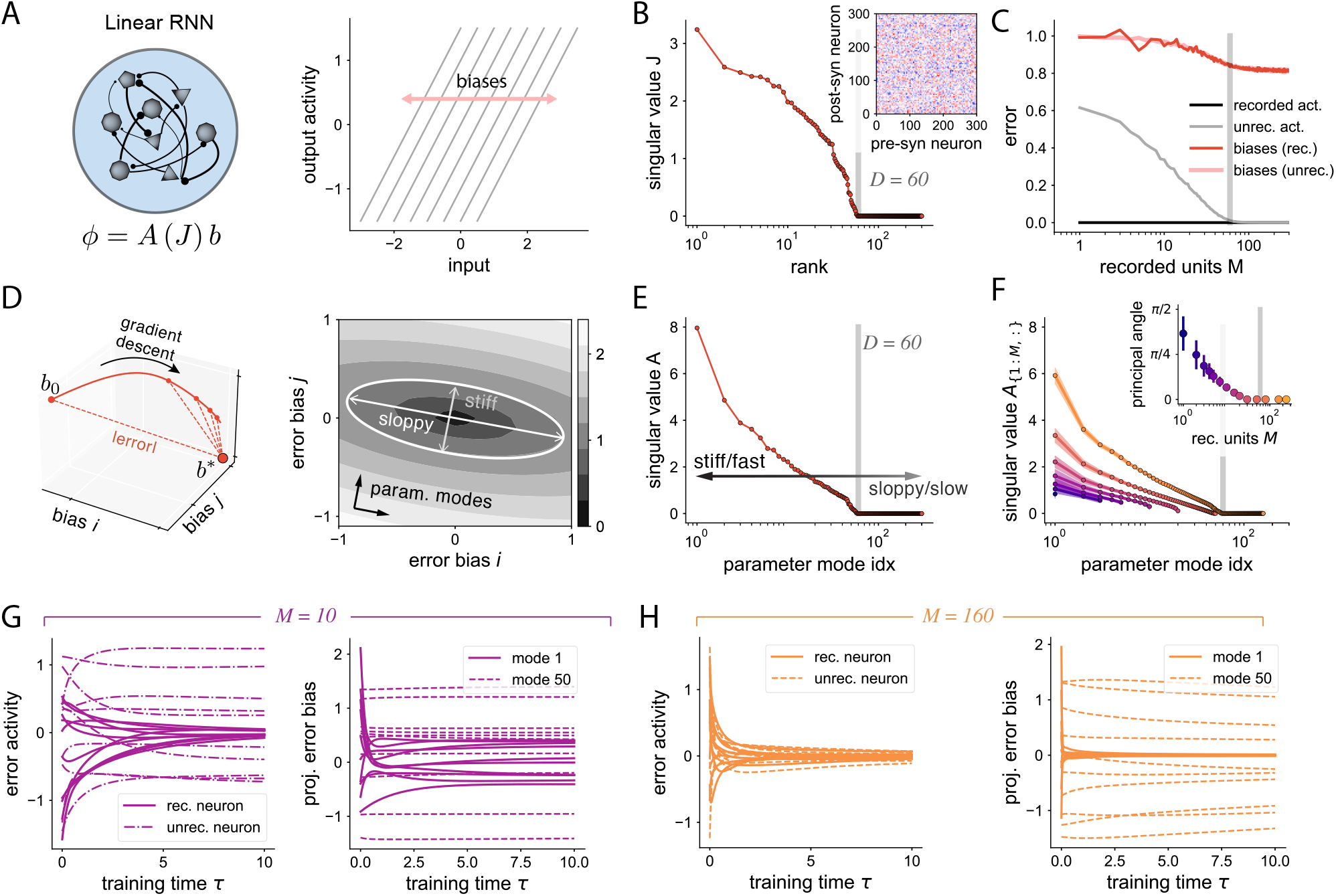
Linear teacher-student model. **A** Left: The activity of a neuron is a linear mapping *A*(*J*), which depends on the connectivity matrix *J*, of the single neuron parameters *b*. Right: Neuronal activation functions are linear with heterogeneous biases. **B** Singular values of the connectivity matrix, which is random and has rank *D* = 60. **C** Errors in activity and biases as a function of the number of recorded neurons. **D** Single neuron biases evolve over training through gradient descent. Parameter modes are described as stiff or sloppy based on the effect of changes along each mode near the optimal solution. **E** Singular value decomposition of the mapping *A* determines stiff and sloppy parameter modes. Stiffer modes are learned more quickly. **F** Effective singular value decomposition when recording from a subset of *M* neurons. Inset shows the maximum angle between the *M* stiffest modes and the *M* sub-sampled parameter modes. **G-H** Evolution of errors in activity and biases for *D > M* = 10 and *D < M* = 160, for 10 different initializations of parameters. Error in biases is projected along one stiff (1st) and one sloppy (50th) parameter mode.

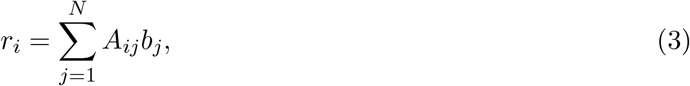

where we have defined *A* ≡ (*I* − *J*)^*†*^ *J*. Although, for simplicity, we focus here on equilibrium activity, time-dependent trajectories would also yield a linear relation between activity and single neuron parameters (see Methods for the full time-dependent derivation). For the same reason, we have also assumed no external input to each neuron (*I*_*i*_ (*t*) = 0). This linear relation between neuronal parameters and activity, which underpins the mathematical tractability of the simplified model, is a consequence of the linear network dynamics and the additive influence of the bias parameters. Choosing multiplicative gains as the unknown single neuron parameters, for instance, would produce a nonlinear relation, which is why we omitted them in this section.

The student is trained using gradient descent on the single neuron parameters. The learning trajectory in parameter space can be expressed, in the limit of small learning rate *η*, as:

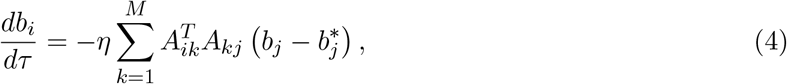

where *M* is the number of recorded neurons. Using these learning dynamics, and because of the explicit mapping between parameters and activity in the simplified model, we can analytically calculate the expected error in recorded and unrecorded activity (see Methods). We show in Fig. 5 C the asymptotic error, after a large number of training epochs, in recorded activity, unrecorded activity, and single neuron biases, for different values of *M*. There is a transition to zero error in the activity of unrecorded neurons when the number of recorded neurons exceeds *D*, the rank of the connectivity matrix (Fig. 5 C, gray line). There are, however, large errors in single neuron parameters (Fig. 5 C, red line) even when the activity of the full network is fully recovered.

To understand these effects, we analyzed the properties of the loss function, which describes how the difference between teacher and student neural activity depends on parameters. We differentiate the loss function for the full network, which is determined by errors in both recorded and unrecorded neural activity, from the loss function for the recorded neurons, which is what is optimized during training. The loss function for the full network is convex with one global minimum (Fig. 5 D). A geometrical analysis demonstrates that the global minimum is surrounded by a valley-shaped region of low loss (Fig. 5D right). We refer to directions for which the loss changes quickly or slowly as the parameters are changed as “stiff” and “sloppy” parameter modes, respectively ^40^. Stiff modes both have the greatest effect on the loss and are learned most quickly. Each mode’s degree of stiffness is determined by the singular values of the matrix *A* (see Methods). When *D < N*, there are *N* − *D* modes with zero singular value, meaning that they do not affect neural activity and are not affected by gradient descent. The presence of these modes explains why single neuron parameters are never fully recovered even when the activity of many neurons is recorded (Fig. 2 C).

The parameter modes that affect the recorded activities and thus determine the loss function for the *M* recorded neurons are specified by *A*_1:*M*,:_ (the submatrix of *A* containing the rows corresponding to these neurons), whose stiff and sloppy modes are generally different from those of the fully sampled matrix *A* (Fig. 5 E vs. F). Subsampling leads to overall smaller singular values, which slows learning dynamics. It also introduces additional modes with zero singular value when *M < D*, since *A*_1:*M*,:_ has at most *M* non-zero singular values. Parameter modes will also, typically, not be fully aligned with those of the fully sampled system (Fig. 5 F, inset). These observations explain why the number of neurons required to achieve a low error depends on *D* (Fig. 2 C, Fig. 3).

We plotted the error in activity and single neuron parameters for a number of recorded neurons below and above the critical number *D* (Fig. 5 G, H). When there are few recorded neurons, the error in recorded neurons eventually goes to zero, while the error in unrecorded neurons decreases on average, but only slightly (Fig. 5 G left). We then examined the error in parameter space. Along the stiffest mode, the error quickly converges to a small value (Fig. 5 G right), while the error along a sloppy mode barely changes. This is because the stiffest mode of the subsampled network is not the stiffest mode of the fully sampled network. When more neurons are recorded, the error in activity decreases toward zero at a similar rate for both recorded and unrecorded neurons (Fig. 5 H left), while the error in parameter space along sloppy modes remains barely affected (Fig. 5 H right).

The simplified model demonstrates that specific patterns of single neuron parameters determine the error between teacher and student. Stiff parameter modes are those that are learned through gradient descent, while sloppy modes are not. When enough neurons are sampled, as compared to the dimensionality of the mapping between parameters and activity, the stiff modes for the *M* recorded neurons align with the stiff modes for the full network, leading to correct prediction of unrecorded activity.

### Loss landscape

We next generalized our theory to nonlinear networks. To facilitate analysis, we studied a class of lowrank RNNs whose activity can be understood analytically ^36;41;42^. We focused on a teacher network with *N* = 1000 neurons and a nonlinear, bounded activation function. We designed the network’s connectivity to be rank-two, with two different subpopulations. Crucially, for this network, there are only two stiff parameter modes, the average single-neuron gain for each subpopulation (Fig. 6 A). We set the connectivity to generate different nonzero fixed points, and we recorded activity as the neural dynamics approached one of these fixed points (Fig. 6 B).

**Figure 6:**
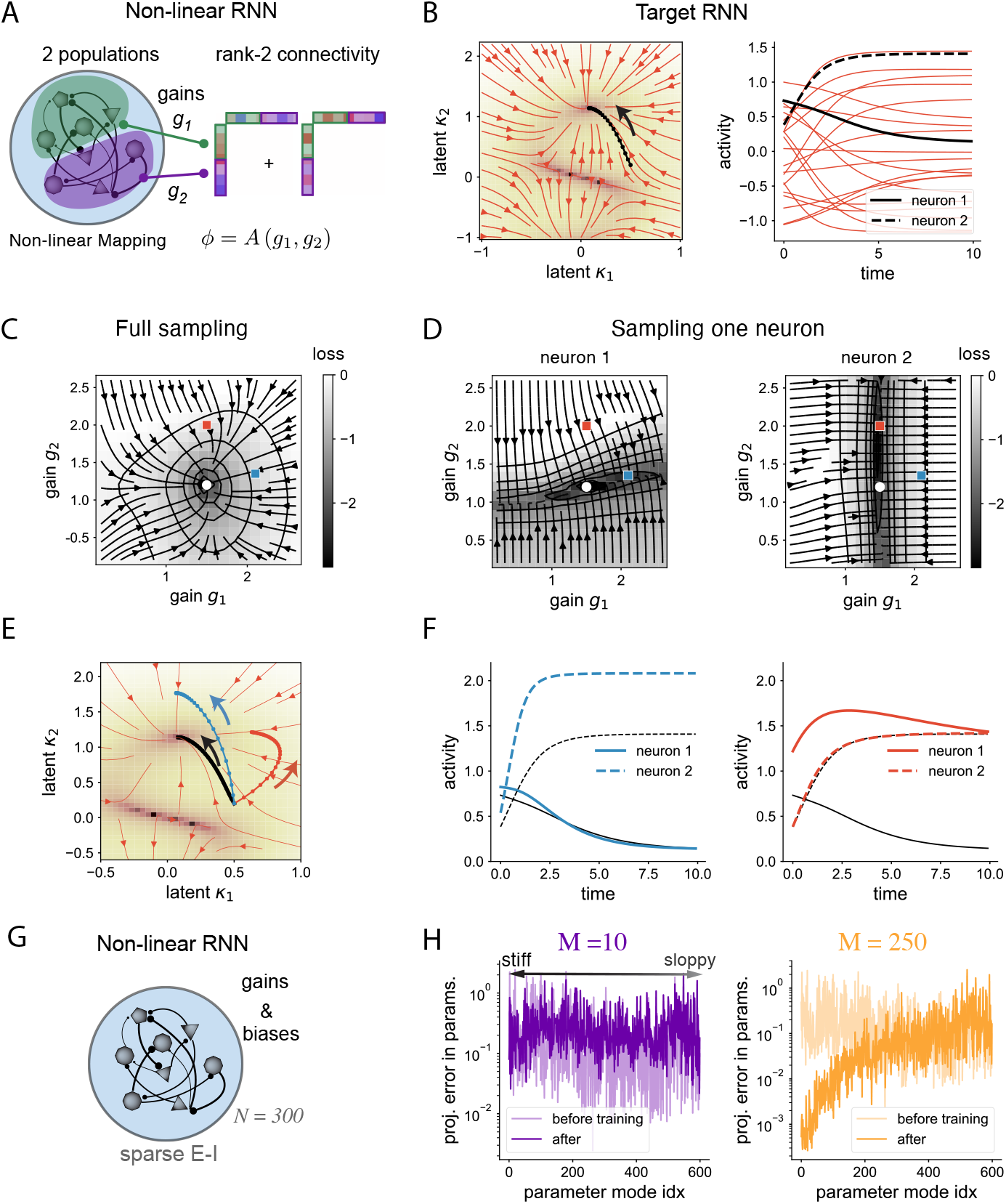
Loss landscape in nonlinear networks. **A** We study a rank-two RNN with two populations. Neurons in each population share the same gains and network statistics. **B** Dynamics of the target network. Left: Phase-space in the two-dimensional latent space. Right: Activity as a function of time for 20 sampled neurons. For illustrative purposes, neurons 1 and 2 are selected based on their alignment with the two latent variables. **C** The loss landscape of the full network depends only on the gains of each population, *g*_1_ and *g*_2_. White dot indicates the parameters of the teacher RNN. **D** Loss landscape when recording the activity of neuron 1 (left) or neuron 2 (right). Blue and red squares correspond respectively to solutions where the training loss is close to zero. **E** Target trajectory (black) and dynamics of the teacher RNN. Blue and red trajectories correspond to the solutions found in D. **F** Predicted activity for neurons 1 and 2 for the solutions found in D. Left: Error in the activity of the recorded neuron (neuron 1) is small, while error for the unrecorded neuron (neuron 2) is large. Right: Similar to left, but when neuron 2 is recorded and neuron 1 is unrecorded. **G** Full-rank non-linear RNN, same as in Fig. 1. **H** Average squared error in parameters projected on the different stiff and sloppy parameter modes. The stiff and sloppy dimensions are determined by approximating the full-sampled loss function around the teacher’s values (see Methods). Average over 10 realizations.

Since the parameter space is two-dimensional, we can visualize the loss landscape for the full network across a grid of parameters (Fig. 6 C). The loss has a single minimum, similar to the linear model. However, due to the nonlinearity, the loss function is non-convex (contour lines are not convex in Fig. 6 C), and the curvature of the loss for parameter values away from the global minimum is different than at the minimum. Despite this non-convexity, gradient descent on this fully sampled loss function will still approach the single minimum.

We next visualized the loss functions for only the recorded neural activity, when activity from one neuron is recorded (Fig. 3 D). Each sampled neuron exhibits distinct dynamics (black lines, Fig. 3 B). The values of the loss function for a single sampled neuron are smaller than for the fully sampled loss, and there is an additional sloppy mode that is not present in the fully sampled loss (black valleys in Fig. 3 D). These results are similar to those of the linear case, although due to the nonlinearity, the sloppy modes correspond to curved regions in parameter space.

The sloppy mode is different for the two recorded neurons. When running gradient descent on these subsampled loss functions, parameter values may therefore move toward the blue dot when neuron 1 is sampled and the red dot when neuron 2 is sampled, since both of these parameter values correspond to a small error in the optimized loss (Fig. 3 D, left and right respectively). However, both of these two solutions also produce high error in unrecorded activity (Fig. 5 C). Indeed, the dynamics of unrecorded neurons for these two solutions deviate substantially from those of the teacher (Fig. 5 E,F).

To test whether the same insights apply also to nonlinear and high-dimensional parameter spaces, we computed the sloppy and stiff modes of the fully sampled loss function in the network of Figs. 1 And 2. We approximated the loss function in parameter space to second order at the optimal parameters. We then projected the average error in parameter space, before and after training, along the estimated stiff and sloppy modes (Fig. 6 G-H). We found that when few neurons are recorded (*M* =10), the average changes in parameter space before and after training are not aligned with the stiff modes of the loss landscape. However, when recording from many neurons (Fig. 6 G-H), there is a large decrease in parameter space along the estimated stiff modes, as predicted by our theory.

We conclude that the qualitative behavior of the linear model holds for nonlinear models. Namely, when the loss function is determined by recordings of a small number of neurons, the parameter modes become sloppier on average, and new sloppy parameter modes are added that do not align with the sloppy modes of the fully sampled loss function.

### Optimal selection of single neurons

So far, the recorded neurons have been selected randomly from the teacher network. As we have seen, different sets of recorded neurons define different loss functions and gradient descent dynamics, suggesting the possibility of selecting recorded neurons to minimize the expected error in unrecorded activity (Fig. 7 A). Specifically, we aim to select recorded neurons such that the sloppy modes of the subsampled loss function align as much as possible with those of the fully sampled loss function.

**Figure 7:**
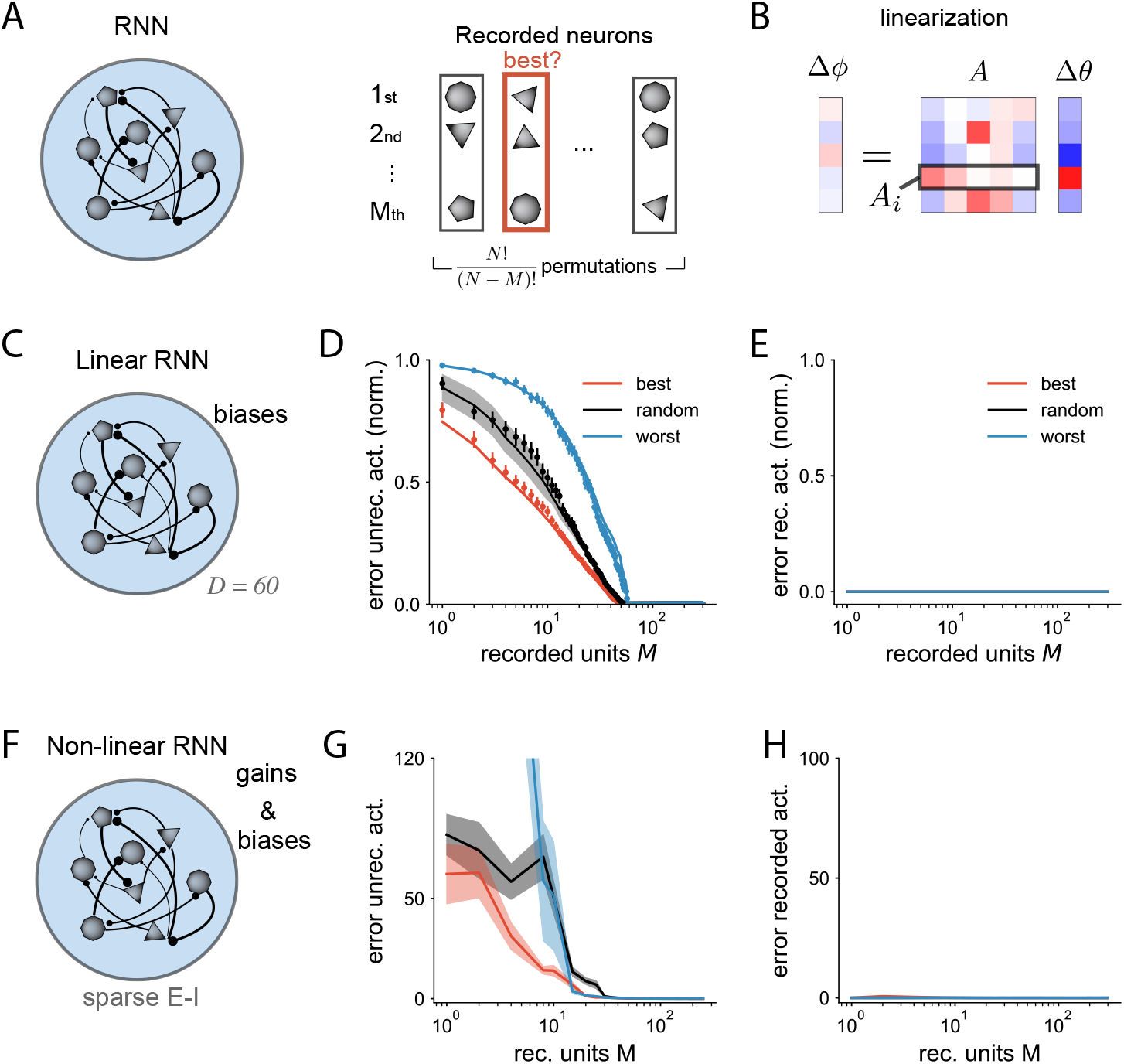
Optimal selection of recorded neurons. **A** Recording from specific subsets of neurons (right) in the teacher RNN leads to different performance. **B** We linearized the mapping from changes in single neuron parameters to changes in neural activity. **C** Teacher RNN with linear single-neuron activation functions, unknown biases, and connectivity with rank *D* = 60 (as in Fig. 5). **D** Error in activity of unrecorded neurons as a function of number of recorded neurons *M*. Lines correspond to theoretical prediction, dots to numerical simulation (mean *±* SEM). We selected neurons following the estimated best ranking (red), 5 different random rankings (black), and the worst ranking (blue). **E** Error in recorded neurons for the same networks. **F - H** Analogous to **C-E** but for a nonlinear network. The teacher is the RNN from Fig. 2 with sparse E-I connectivity. Single-neuron parameters are both gains and biases. The linearization in the mapping from parameters to activity assumes homogeneous single-neuron parameters (see Methods).

In the simplified linear model, subsampling neurons corresponds to selecting rows of the matrix *A* that relates single neuron parameters to activity (Fig. 7 B). In this case, it is possible to exactly determine which neurons are most informative to record. Formally, the most informative neuron *i* is the one whose corresponding row *A*_*i*,:_ overlaps most with the weighted left singular vectors of the matrix *A* (see Methods). The second most informative neuron is the one whose row overlaps most with the weighted left singular vectors of *A* that are orthogonal to the previously selected neurons, and so on. It is also possible to define the least informative sequence of recorded neurons by minimizing, rather than maximizing, these overlaps. We compared the error in unrecorded activity for the most and least informative sequence of selected neurons as well as random selection, finding that the optimal strategy indeed improves the efficiency of training (Fig. 7 D).

For nonlinear networks, the mapping between parameters and network activity is also nonlinear and depends on the unknown parameters of the teacher (see Methods). As a result, the globally optimal sequence of neurons to record from cannot be determined a priori. Nevertheless, the mapping between parameters and activity can be linearized based on an initial guess of the single neuron parameters and then iteratively refined. In practice, we found that linearization works well for nonlinear networks, with the optimal selection strategy dramatically reducing the error compared to random selection. For the network studied in Fig. 2, the error using the best 10 predicted neurons is 60% smaller than random selection (Fig. 7 G).

The singular vectors used to determine which neurons are most informative depend on the global connectivity structure and cannot be exactly reduced to any single neuron property. Such properties, including in-degree, out-degree, average synaptic strength, or neuron firing rate, may be correlated with the singular value decomposition score developed here, but are not guaranteed to be good proxies for informativeness. These results thus argue for the use of models like those we have studied here to guide the selection of recorded neurons.

## DISCUSSION

Building connectivity-constrained neural network models has become increasingly viable as the scale of connectome datasets has grown. Our theory cautions against against over-interpreting such models when they are insufficiently constrained (Fig. 1), but also shows that correctly-parameterized models paired with sufficiently many neural recordings can provide consistent predictions (Fig. 2). This consistency is a consequence of the qualitatively different solution spaces associated with connectome-constrained and unconstrained models (Fig. 2C,D, Fig. 6). The theory also suggests that models can be used to inform targets for physiological recordings (Fig. 7).

### Challenges for connectome-constrained neural networks

We have studied the properties of connectome-constrained neural networks using simulations, as aligned physiological and connectomic datasets are not widely available. Our results suggest that the “forward problem” of predicting neural activity using a connectome is not as ill-posed as the corresponding “inverse problem” studied previously ^1^. However, although we demonstrated that this result is robust to model mismatch and inaccuracy in synaptic reconstruction (Fig. 4), it is likely that for some neural systems the degree of model mismatch is too severe. Such systems likely include those for which the dynamics are largely driven by unmodeled processes such as the effects of neuropeptides or gap junction coupling^13;23^. Moreover, systems for which the firing rate models described here are a poor match, such as systems that operate based on spike synchrony rather than rate codes^43^, highly compartmentalized interactions^44^, or dynamics of specific ion channels^45^, may be out of reach of the present approach. We have also assumed that time-varying external inputs to the network are known, and we do not expect connectomes to provide substantial constraints on strongly input-driven neural activity when inputs are not controlled. Nonetheless, our results establish for the first time that, appropriately, the dynamics in RNNs with order *N*, rather than *N* ^2^, unknown parameters, can be accurately predicted, unlike the corresponding inverse problem.

We note that the parameters in our models may involve state-dependent modulation. Neuromodulators, for instance, are known to modify effective neuronal excitabilities^27^. In our networks, gains and biases do not necessarily account for a single biophysical process but rather the coordinated effects of multiple processes. As long as the timescale of these processes is slower than the dynamics being predicted, we expect an approach similar to the one described here to be appropriate. However, this state-dependence may also imply that the inferred parameters may not generalize to new behavioral states.

### Assessment of connectome-constrained solutions

It is known that, in order to estimate neural dynamics lying in a manifold of linear dimensionality *D*, it is necessary to record from *N > D* different neurons, independent of network size ^33^. Connectome-constrained models go beyond such mean-field or population-level descriptions of neural dynamics, as they are also concerned with how each specific neuron contributes to global activity patterns. This requires knowledge of unrecorded neurons’ loadings onto the low-dimensional manifold. This benchmark is appropriate when such models are used to predict the function of specific neurons or neuron types, or to guide experiments that manipulate specific neurons^14;16;19^.

The match between student and teacher activities depends on multiple properties. These include whether random choices of single neuron parameters produce similar dynamics in the two networks (Fig. 3C, Supp. Fig. 9), the extent of model mismatch (Fig. 4), and the degree to which training the student based on activity recordings reduces this error (Supp. Fig. 8). The first two of these depend on specific features of the teacher network. Recent studies have demonstrated above-chance prediction of function using uniform or random parameters in models of the *Drosophila* nervous system^14;19^. These studies focused on largely feedforward circuits in which precise values of single neuron parameters may not be required for accurate predictions. On the other hand, failure of a related approach in *C. elegans* was argued to be a consequence of model mismatch from unmodeled peptidergic interactions^13^.

We found that training student networks to match the teacher only led to improvements when, in addition to connectivity constraints, sufficiently many neurons’ activities were constrained. This result was independent of the initial performance of the system with random parameters (Supp. Fig. 9). It is possible that, for networks that perform multiple tasks or process diverse inputs, recording activity under multiple task or input conditions may lead to a similar improvement, if single neuron parameters are not strongly modulated across these conditions.

### Properties of connectome-constrained and unconstrained network solutions

The loss function of feedforward neural networks trained on different tasks has been shown to have multiple minima, with often counter-intuitive geometrical properties ^46;47^. The multiplicity of minima arises from symmetries such as weight permutations in the network parametrization^48^. It remains unclear whether such ideas extend to recurrent neural networks. We found that for connectivity-constrained networks in a teacher-student paradigm, our results are consistent with the existence of a single minimum with stiff and sloppy parameter modes around the optimal solution (Fig. 6). The alignment between sloppy parameter modes in subsampled versus fully sampled loss functions explains the success in generalizing to unrecorded neurons.

Robustness to a large range of structural parameters and perturbations is a hallmark of biological systems, with a few stiff parameter combinations determining function^40;49–51^. We have shown that this is also true of connectome-constrained networks. One consequence of this observation is that, in datadriven models for neuroscience and machine learning that are largely underconstrained, the distribution of parameters such as synaptic weights or single neuron excitabilities found after successful training may not be predictive of task performance. Our work argues in favor of identifying stiff parameter combinations in such networks and using these to assess the similarity of network solutions ^52^.

## ACKNOWLEDGEMENTS

The authors are grateful to L.F. Abbott for helpful discussions and comments on a previous version of the manuscript. M.B. and A.L.-K were supported by NIH award R01EB029858 and the Gatsby Charitable Foundation GAT3708. A.L.-K. was supported by the McKnight Endowment Fund.

## CODE AVAILABILITY

Code and trained networks will be made available upon publication.

## METHODS

### Recurrent network models

We focused on recurrent neural networks where the activity of each neuron *i* is described by a continuous variable, a firing rate *r*_*i*_ (*t*), for *i* = 1 … *N*. The firing rate of each neuron is calculated by applying the a parametric function to the input *x*_*i*_ (*t*) that the neuron receives at each time point,

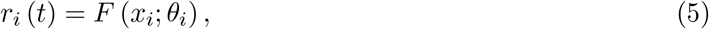

where *θ*_*i*_ denote the single neuron parameters that modulate the function *F*. We denote this input-to-rate function *F* as activation function or input-output function. The activation function may depend on different single neuron parameters *θ*_*i*_, such as the gains *g*_*i*_ and biases *b*_*i*_, as shown in Eq. (1).

The dynamics of the recurrent neural network is defined at the level of the inputs as

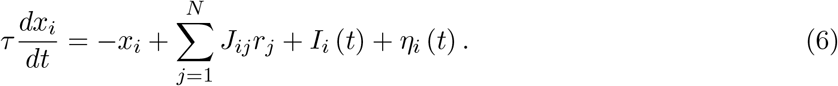

The matrix **J** is the synaptic connectivity matrix, the element *J*_*ij*_ indicated the signed synaptic strength of the connection from neuron *j* to neuron *i*. We explicitly separated the external input into the term *I*_*i*_ (*t*), representing the task related external inputs, and the private white noise given to each neuron *η*_*i*_ (*t*) which is added to provide stability in the solutions. We set the single neuron time constant to unity, and expressed all temporal quantities with respect to this time constant.

The dynamical landscape that a network can implement is thus determined by the order *N* single neuron parameters and the *N* ^2^ synaptic strengths of the connectivity. In the section Teacher RNNs below, we specify the choice of activation function, single neuron parameters and synaptic connectivity matrices used in each figure.

### Teacher-student: setup and training

We focus on a set of two RNNs: one of them is the teacher RNN, which represents the network from which we know the connectivity and from which we can record neural activity. The other network is the student RNN, which is trained -its parameters are optimizedto match neural activity in the teacher. Both networks’ dynamics are determined by Eq. (6). We use the asterisk notation on network parameters when we refer specifically to the parameters in the teacher’s network.

Unless otherwise specified, the teacher and the student network share the same connectivity matrix **J**. Additionally, in all cases, the external input *I*_*i*_ (*t*) for all neurons *i* = 1 … *N* and times *t* and the initial conditions *x*_*i*_ (*t* = 0) are the same in teacher and student. The possible structural differences between teacher and student lie in the set of the single neuron parameters *{θ*_*i*_*}*. These single neuron parameters are optimized to match the recorded activity of the teacher. In Fig. 2 D and Supplementary Fig. 8, where we train students with unknown connectivity, we set the single neuron parameters to be equal in teacher and student. The connectivity matrices *J* are different in teacher and student, and the synaptic weights are trained in the student.

The trained parameters are optimized to minimize the loss function

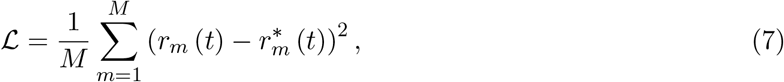

where the square brackets denote average over the time points in the recorded window, and we sorted neurons such that the first *M* neurons are the recorded ones. In Fig. 1, instead of defining the loss on the recorded activity *r*_*m*_ (*t*), we used on the task readout, 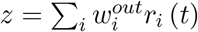.

We trained the parameters of the student using standard gradient descent methods applied to time varying signals: we implemented backpropagation through time via the ADAM optimized using pytorch^53–55^. We used learning rates varying between 0.0001 and 0.01, and decay rates of the first and second moments of 0.9 and 0.999.

### Quantifying performance

We used mean-squared error as the measure for quantifying deviations between teacher and student, unless otherwise specified. In particular, we assessed three different types of errors: error in the activity of recorded neurons, error in the activity of unrecorded neurons and error between trainable parameters.

In Fig. 3, we quantified the activity error using the Pearson correlation coefficient of each neuron *ρ*_*i*_:

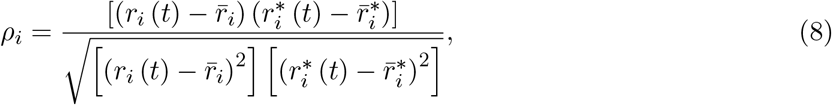

where the square brackets indicate average across timepoints and 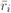 is the average activity of the *i*-th neuron. The error is then defined as the Pearson’s distance, one minus the absolute value of the Pearson correlation coefficient averaged across recorded and unrecorded neurons. This measure of dissimilarity allows us to measure whether the temporal profile of teacher and student are similar or not, even if there is a mismatch in the mean and scaling factors. Large mismatches in mean and amplitude will dominate the mean squared error, even when the temporal dynamics are well reproduced by the student, due to the design of the teacher network.

In Fig. 2, F or matching the activity and connectivity between unrecorded neurons, we paired unrecorded neurons in the teacher with unrecorded neurons in the student by following a greedy procedure, where we selected random neurons in the teacher and searched for the most similar neuron in terms of activity in the student. Once a pairing is done, both neurons in the teacher and student are removed, and a new teacher candidate is selected.

### Teacher RNNs and training parameters

In this next section, we detail the choice of single-neuron parameters, the features of the teacher RNN, and the training parameters used in the different figures. Unless otherwise specified, we used Δ*t* = 0.1 in units of the single-neuron time constant. We injected noise at each timestep of the dynamics with standard deviation 0.002. The initial guesses for the unknown parameters (usually the single neuron gains and/or biases) are random permutations of the teacher’s parameters. This assumes that the mean and variance of the single neuron parameters are known. See also shared code, for reproducing all the numerical experiments in the study.

**Figures 1 And 2:** The teacher RNN (*N* = 300) is trained with learning rate 0.001 during 1400 epochs. The initial connectivity is chosen as follows: first, each synapse is drawn from a random Gaussian distribution with mean zero and variance 2.4/np.sqrt(N). The sparse weights (fraction *p* = 0.5) are randomly chosen, set to zero, and not trained. A fraction *p*_*E*_ = 0.7 of neurons selected randomly is set to be excitatory, such that their synaptic strengths are set to their absolute value, while the remaining fraction of neurons, 1 − *p*_*E*_, is set to be inhibitory. Synapses are rectified to their assigned sign after each training epoch. Trials are 20 time units long. The single neuron activation function is given by Softplus:

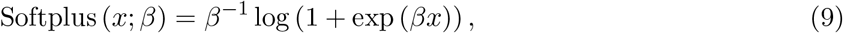

where we set the smoothness parameter to *beta* = 1.

The student RNNs with unknown single neuron parameters are trained for 7000 epochs and learning rate 0.001. The student RNN with unknown connectivity shares the same single neuron parameters as the teacher RNN, to facilitate the comparison. The figure shows results for 14 trained different students. Learning rate: 0.005. The synaptic weights are initialized randomly following a Gaussian distribution, and the weight signs are correctly assigned (i.e., the student knows whether a neuron is excitatory or inhibitory). To compare the weights after training, we picked a random unrecorded neuron from the teacher and matched it with the unrecorded neuron with the most similar activity profile. Then, the selected neurons in teacher and student are discarded, and we picked a new neuron to be matched in the teacher. This procedure is repeated until all neurons are paired.

**Figure 3:** The teacher networks in the top row are rank-two networks with the same correlations across the singular vectors of the connectivity, such that in the limit of large *N*, networks converge to the same mean-field dynamics. The activation function for each neuron is tanh, and we consider gains as the only single neuron parameter. The connectivity loadings from each neuron are sampled from a multivariate Gaussian distribution with zero-mean. The loadings of left singular vectors have all unit variance, while the loadings of right singular vectors have the same variance, large enough such that the covariance of all connectivity entries is positive definite. Following the notation in ^36^, we chose 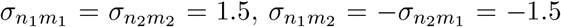, such that the network generates a stable limit cycle. The gains are chosen to be Gaussian, with mean 1 and standard deviation 0.9, uncorrelated with all the other connectivity loadings. We selected trajectories that start on the limit cycle, and evolve during 20 time units. The student network is initialized in this case with homogeneous unitary gains, and is trained for 7000 epochs with learning rate 0.005.

The teacher networks in the bottom row have random connectivity as in^37^, where the synaptic strengths *J*_*ij*_ are randomly drawn from a Gaussian distribution with mean zero and standard deviation 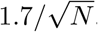. The single-neuron gains are drawn from a Gaussian distribution of unit mean and standard deviation 0.5.

The student networks were initialized before training with homogeneous gains, *g*_*i*_ = 1.

**Figure 4:** The teacher network is the same biologically-inspired network as in Fig. 2.

**Figure 5:** For the connectivity of the teacher in Fig. 5 B-H, we drew each synaptic strength *J*_*ij*_ from a Gaussian distribution with mean zero and standard deviation 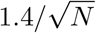, and based on the singular value decomposition, kept the first 60 rank-one components. Network size *N* = 300.

In Fig. 5F (inset), to calculate the principle angle between the first *M* singular vectors of the subsampled matrix *A*_1:*M*,:_ and the full matrix *A*, we calculated the *M* left singular vectors of the matrix 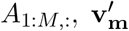, and the first *M* left singular vectors of the matrix *A*, **v**_**m**_. The principle angle measures the maximum angle between two linear subspaces. We computed the principal angle as the maximum singular value of the matrix product 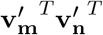, for *m, n* = 1 … *M*.

**Figure 6:** We designed a teacher RNN with rank-two connectivity and two populations^36^, tanh activation function and gains as single neuron parameters, *N* = 1000 neurons. The parameters in the first population are: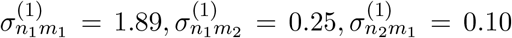 and 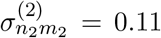 in the second population 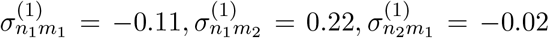, and 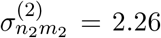. The gains are reduced to a two-dimensional parameter space, where all the gains of neurons in population 1 have the same value, *g*_1_, and all the gains of neurons in population 2 have valu e *g*_2_. In the teacher network, 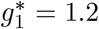 and 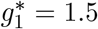. The parameters are chosen such that the first population has more control of the dynamics along the variable *κ*_1_, and the second population controls *κ*_2_.

**Figure 7:** The linear network corresponds to the same network as in Fig. 5. The non-linear network is the same network as in Fig. 2.

### Prediction in linear recurrent networks

In the linear model, the RNNs are linear networks with dynamics

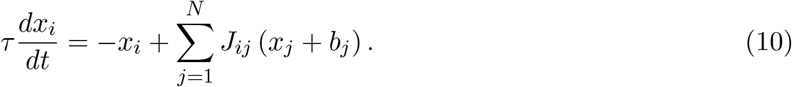

We define the activity in this linear network as *r*_*i*_ (*t*) = *x*_*i*_ (*t*). The single-neuron parameters *b*_*i*_ correspond to the bias. Throughout the results section, we focused on the fixed point activity, which is given in vector form by

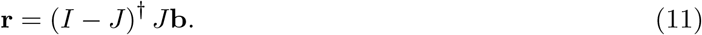

where *I* is the identity matrix. There is a linear mapping between single neuron parameters **b** and activity **r**, given my a matrix *A*, in this case defined as *A* = (*I* − *J*)^*†*^ *J*. The notation *A*^*†*^ indicates the pseudo-inverse.

### Fully sampled teacher

The loss function when all neurons are recorded, is given by the quadratic form

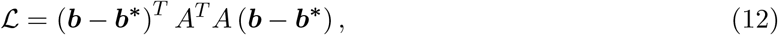

such that there is one global minimum when the student and teacher are identical to each other, ***b*** = ***b***^**∗**^, and the Hessian of the loss is independent of the teacher’s biases ***b***^**∗**^. Running gradient descent, in the limit of small learning rates *η*, leads to the following equation for the estimated biases in the student over the timecourse *τ* of learning:

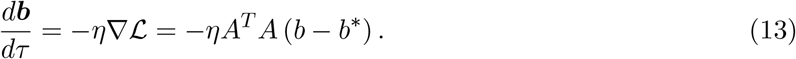

The singular value decomposition of matrix 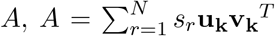, determines the training dynamics. We refer to the left singular vector **u**_**k**_ as an activity mode, and right singular vector **v**_**k**_, a parameter mode. The error in parameter space along mode **v**_**k**_ decreases over training time with timescale given by 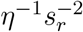, reducing the error in activity along the mode **u**_**k**_. An initial guess **b**_**0**_ which is a distance of one away from the teacher **b**^∗^ along mode **v**_**k**_ generates an error in the activity of neurons along mode **u**_**k**_ and magnitude *s*_*k*_. Thus, parameter modes that produce big errors in the activity are learned fast, while parameter modes that produce small errors in the activity are learned more slowly. We refer to parameter modes corresponding to large singular values as *stiff* modes, and *sloppy* modes as those parameter modes with very low singular value.

If the connectivity *J* is not full-rank, some singular values of the mapping matrix *A* will be zero. In that case, the parameter values along the modes *v*_*k*_ corresponding to singular value *s*_*k*_ = 0 (the extreme case of sloppy parameter modes) cannot be inferred through gradient descent, although that mismatch does not cause any error in the activity of unrecorded neurons.

All the results can be directly extended to linear networks where transient trajectories are considered, given an initial state **x**_**0**_. The considered dynamics would read

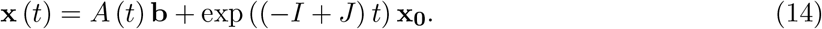

where there is an affine mapping from **x** (*t*) to parameters **b**, now given by the tensor *A* (*t*):

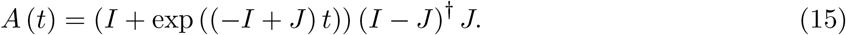

In the case of a temporally dependent mapping *A* (*t*), the relevant singular values are the square root of the eigenvalues of the time-averaged matrix [*A* (*t*) *A* (*t*)^*T*^], and the activity modes are its eigenvectors.

The parameter modes are the eigenvectors of the time-averaged matrix [*A* (*t*)^*T*^ *A* (*t*)].

### Subsampled activity

Recording from a subsample of *M* neurons is equivalent to selecting the rows of matrix *A* corresponding to the recorded neurons, and removing the rest. We refer to this matrix as matrix [*A*]_1:*M*,:_. Equations (12) and (13) still hold, when substituting [*A*]_1:*M*,:_ for *A*.

The effect of subsampling limits the number of learnable or stiff parameter modes of the loss function used for training, which cannot be more than *M*. The fact that the initial guess **b**_**0**_ can only be corrected along *M* modes makes the error in the unrecorded activity be non-zero, when the rank of *A* is larger than *M*, i.e. when not enough neurons are sampled. Furthermore, the parameter modes and activity modes without a non-zero eigenvalue of the training loss need not align with the stiffest modes of the fully sampled loss function.

### One recorded neuron

In linear networks, we can calculate the average error when we record only from neuron *i*. We refer to the subsampled matrix *A*_*i*,:_, which corresponds to a vector as **a**_**i**_. After training a student with initial parameters **b**_**0**_ for *τ* long enough, which is equivalent to assuming a zero-error in the recorded activity of the student, the vector of biases after training, **b**_**f**_ reads

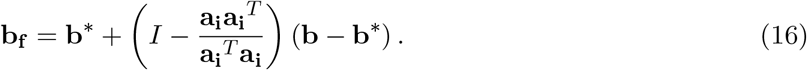

The error in single neuron parameters (combining both recorded and unrecorded neurons), *e*_*f*_, is calculated based on the norm of the vector **b**_**f**_ − **b**_**0**_ given by Eq. 16, which reads:

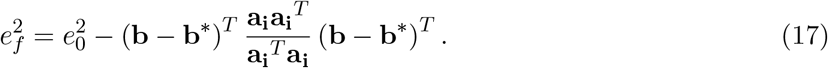

Assuming that the initial guesses **b**_**0**_ are unbiased with respect to the teacher parameters **b**^2^, on average over initial conditions, the improvement in parameter error is

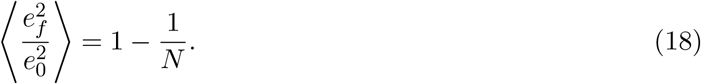

Therefore, on average, the error in parameter space is equally reduced for any selected neuron.

The error in the activity of unrecorded neurons, *E*_*f*_, using the singular value decompositon of *A* is defined as:

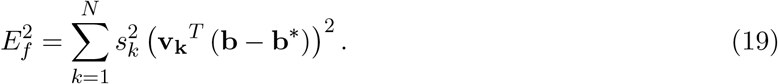

The error in the unrecorded activity 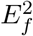 can be larger or smaller than the error before training, unlike the error in parameter space, which can only decrease. Nevertheless, on average over initial conditions, the expected error always decreases and reads

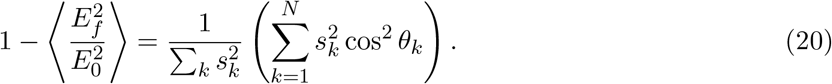

where cos *θ*_*k*_ corresponds to the angle between **a**_**i**_ and **v**_**k**_. Eq. 20 is used to calculate the theoretical predictions in Fig. 7E.

### Optimal selection of neurons: linear RNN

To calculate the best and worst strategy for sampling neurons (Fig. 7) we used a greedy strategy, where we first selected the neuron with the highest and lowest expected reduction in activity error, based on Eq. 20. Then, we proceeded iteratively, projecting out the component 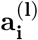 from the (*l*)-th selected neuron from the matrix *A*^(*t*)^, calculating the matrix *A*^(*l*+1)^:

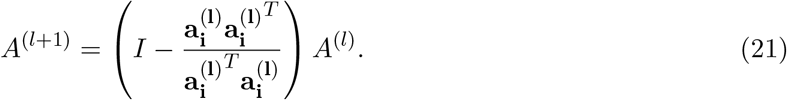

We then selected again the row-vector 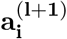 that maximizes (minimizes) the decrease error in Eq. 20, For the best (worst) greedy selection of neurons.

### Optimal selection of neurons: non-linear RNN

For any teacher RNN with unknown gains or non-linear activation functions, the mapping between unknown single neuron parameters and activity is not given by a linear transformation via a matrix *A*. Moreover, the linearization of the gradient dynamics (Eq. 13) close to the teacher parameter depends on the specific parameters, unlike the linear case. Nevertheless, we can still compute the best and worst selection of neurons based on an initial guess of the target parameters.

We focus on activation functions of the form **r** = ĝ *ϕ* (**x** + **b**), where 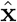 represents a diagonal matrix whose non-zero elements are given by vector **x** and we assume the function *ϕ* is invertible. We are interested in the linearization Δ**r***/*Δ**b** and Δ**r***/*Δ**g**. We focused on fixed point activity, therefore, using Eq. (13), we can define the function:d

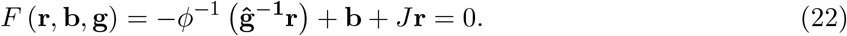

By using the implicit function theorem on *F*, we can calculate the linearized mapping from parameters to activity:

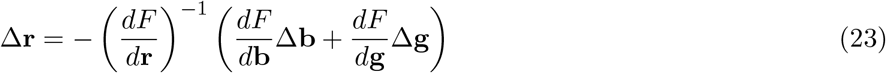

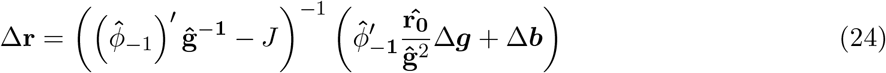

This linear relationships allows us to define a linear matrix *A* when we use random guesses for the teacher parameters, and apply the same algorithm as in linear RNNs. The algorithm for nonlinear networks or gain parameters works by assuming that the curvature of the loss function close to the guessed parameters is similar to the curvature of the loss function close to the teacher parameters.

In Fig. 6 G-H, the Jacobian of the mapping between time-varying activity and single-neuron parameters around the teacher’s parameter values was performed numerically, using Pytorch’s automatic differentiation.

## SUPPLEMENTARY MATERIAL

**Supplementary Figure 1:**
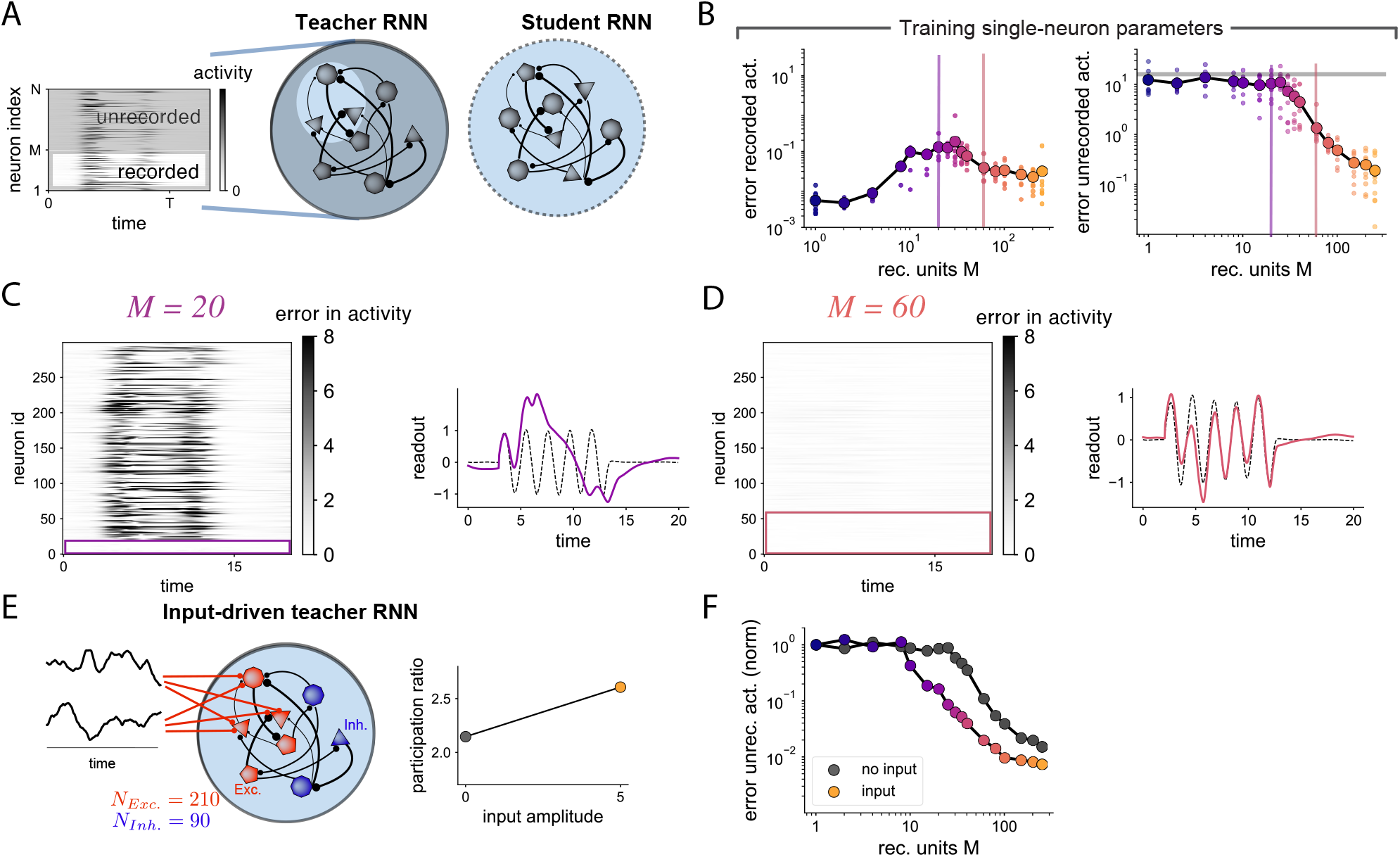
Related to Fig. 2. **A** Teacher as in Fig. 2. The students are trained on a varying number of recorded neurons *M*. **B** Average error in the recorded and unrecorded activity between teacher and students. **C** Left: Error in the network activity for a given student network in a given trial, when *M* = 20 neurons are recorded. Right: Error in the task-related readout signal. While the recorded neurons have low error, the unrecorded neurons in the student display large deviations. **D** Analogous to **C**, when more neurons are recorded, *M* = 60. In this case, the activity of unrecorded neurons and the readout are well predicted. **E** Teacher network receives a strong external two-dimensional time-varying input, fed to a subset of 100 excitatory neurons. Middle: The dimensionality of the activity, measured by the participation ratio, increases with the input. Right: Error in unrecorded neurons after training student networks to match the input-driven teacher (color dots), compared to the non-driven teacher (grey dots). Fewer recorded neurons are required to predict activity of unrecorded neurons in this example input-driven network.

**Supplementary Figure 2:**
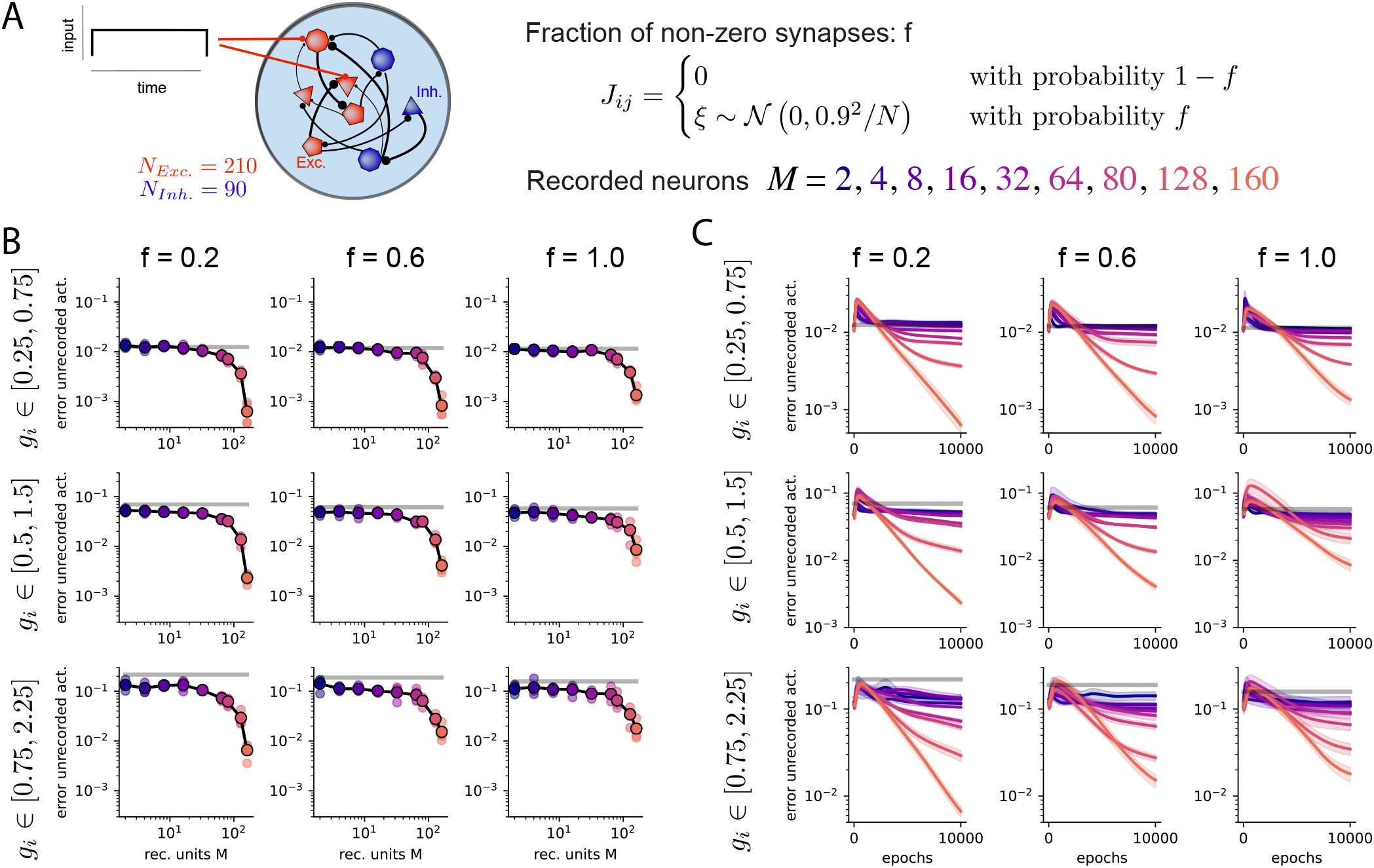
Related to Fig. 2. Input-driven teacher networks with different levels of connectivity sparsity and gain heterogeneity. **A** Teachers have E-I connectivity, and are initialized at the fixed point. A positive input of unit strength is delivered to 5 excitatory neurons. The recorded neurons correspond to excitatory neurons, while unrecorded neurons can be both excitatory or inhibitory. Teacher networks are generated with different fractions *f* of non-zero weights, and different ranges for the uniformly distributed gains. Both gains and biases are trained in the students. **B** Error in unrecorded activity after training vs number of recorded neurons, for different sparsities and gain distributions. While the overall magnitude of the error changes for different gain strengths, the decay of the error as a function of *M* does not change. **C** Evolution of the error in unrecorded activity during training. Note that for high gains, the initial error before training is below baseline.

**Supplementary Figure 3:**
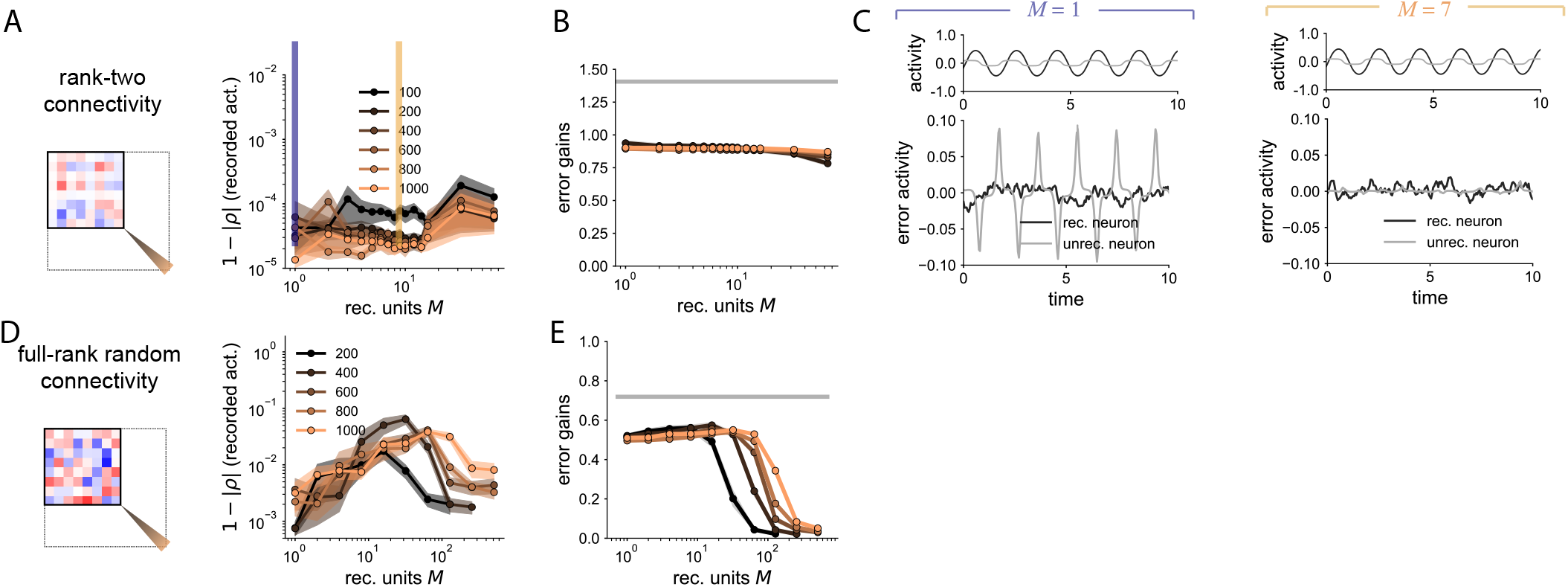
Related to Fig. 3. Teacher networks with different dynamics. **A** Teachers with variable network size and fixed rank-two connectivity, generating a limit cycle. Right: Error in the activity of recorded neurons after training. The students always learn the dynamics of the teacher. **B** Error in the single-neuron gains after training. **C** Example of error in the activity of a recorded neuron and an unrecorded neuron, when there is only one recorded neuron (left), compared to when 7 neurons are recorded (right). For one recorded neuron, the student learns the frequency of the limit cycle, but the temporal profile of the unrecorded neurons does not much the profile of the teacher network. Example for *N* = 400. **D** Teachers with variable network size and random connectivity, generating chaotic dynamics. Right: Error in the activity of recorded neurons after training. The students always learn the dynamics of the teacher. **E** Error in the single neuron gains for the chaotic teachers.

**Supplementary Figure 4:**
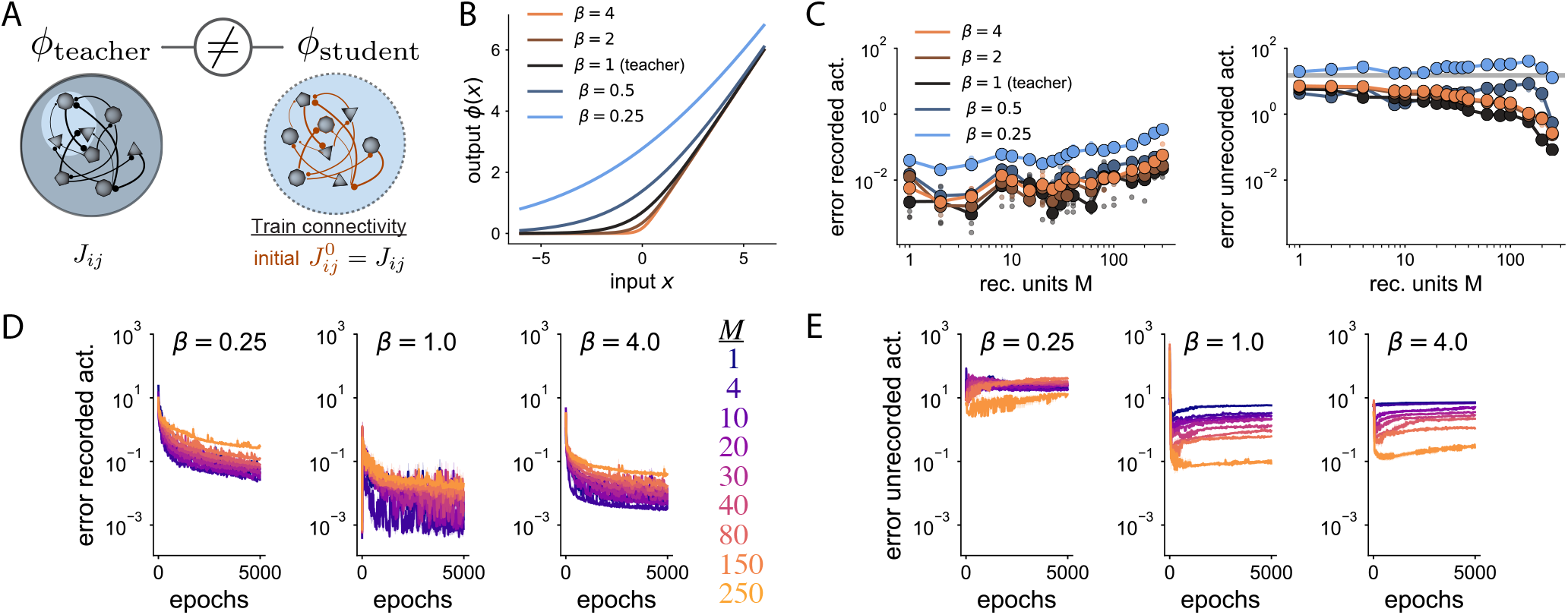
Related to Fig. 4. Training connectivity with model mismatch between teacher and student. **A** We train the connectivity of the student, given the teacher’s connectivity as initial condition. The single neuron parameters are the same in teacher and student, while there is a mismatch in the activation function. Same network as in Fig. 4. **B** The activation function is a smooth rectification but with different degrees of smoothness, parameterized by a parameter *β*. Teacher RNN from Fig. 2. **C** Errors in the activity of recorded (left) and unrecorded (right) neurons for different values of model mismatch between teacher and student. We observe a minor decrease in the error in unrecorded neurons when recording from a large number of neurons, *M* ≈ 150. **D** Error in the recorded activity (loss function) for three different mismatch values as a function of training epochs (*β* = 1. means no mismatch). **E** Error in the unrecorded activity (loss function) for three different mismatch values as a function of training epochs.

